# egtplot: A Python Package for 3-Strategy Evolutionary Games

**DOI:** 10.1101/300004

**Authors:** Inom Mirzaev, Drew FK Williamson, Jacob G Scott

**Affiliations:** Mathematical Biosciences Institute, The Ohio State University; Department of Translational Hematology and Oncology Research, Cleveland Clinic Foundation; Department of Radiation Oncology, Cleveland Clinic Foundation

## 1 Introduction

Evolutionary game theory is a reformulation of classical game theory wherein the players of the game are members of a population. These members do not choose a strategy, but instead are born with their strategy ingrained‐‐they cannot change strategy during the game. In biological terms, the strategies might represent discrete species or genotypes, hawks and doves being a classic example. Payoffs the players gain or lose based on their interactions with other players increase or decrease their fitness, thereby influencing the number or proportion of members playing that strategy in the next generation. As such, the populations of strategies can wax and wane as they outcompete or are outcompeted by other strategies.

As a very general model of cooperation and competition, EGT is well-suited to quantitative investigations of the dynamics of interactions between populations. EGT has been used to model phenomena from disparate areas of study, from poker [1] to hawks and doves [2] to host-parasite coevolution [3]. In many biological EGT models, payoffs are taken to represent the resources a particular organism can extract from its environment given its interaction in that environment with another organism utilizing the same or perhaps a different strategy.

### 1.1 The Replicator Equation

First introduced by Taylor and Jonker [4] in 1978, the replicator equations is one of the most important game dynamics in EGT. In its the most general mathematical form, the replicator equations are given by

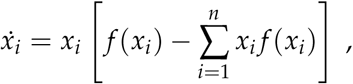

where *X_i_* and *f* (*x_i_*) is the ratio and fitness of type of *i* in the population. The equation is defined on *n*-dimensional simplex and the population vector, *x* = (*x*_1_,…, *x_n_*), sums to unity. In biological terms, per capita change in type *i* (i.e., *ẋ_i_*/*x_i_*) in a well-mixed population is equal to the difference between its expected fitness and the weighted average fitness of the population.

For the sake of simplicity it is often assumed that fitness is linearly proportional to the population distribution. In this case the replicator equations can be written as

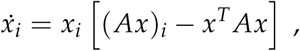

where the matrix A is a payoff matrix with element *A_ij_* representing fitness of type *i* over type *j*.

### 1.2 An Example

Though the replicator equation can model evolutionary games with any natural number of strategies, in this software package, we assume the above form of the replicator equations on a 3-dimensional simplex, meaning we model only three-strategy evolutionary games. Threedimensional replicator equations are well-characterized [5] and can be classified into 46 qualitatively different phase portraits. As an example, we visualize dynamic 34 from [5]:

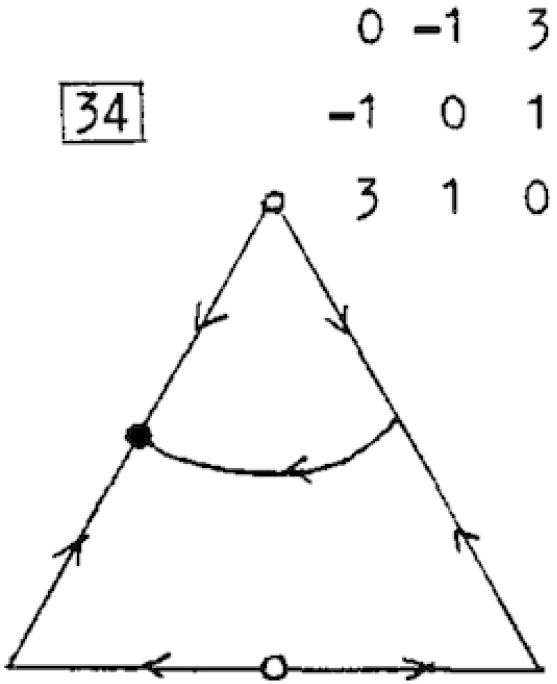

This example shows a stable equilibrium on the left edge, unstable equilibria at the top vertex and on the bottom edge, and a saddle point on the right edge. Let us compare this qualitative plot of this particular evolutionary game to the quantitative plot that our package, `egtplot` can produce. For reference, we use `scipy.integrate.odeint` to solve the replicator equation, which in turn uses Isoda from the FORTRAN library odepack [6].

To demonstrate the functionalities of our software, we begin by importing the package.

~~~
In [1]: from egtplot import plot_static
~~~

The function `plot_static` takes a list of list of parameters with the first three elements holding lists of values for the zeroth row of the payoff matrix, the next three for the first row, and the final three for the second row. For example, the list of lists `[[a], [b], [c], [d], [e], [f], [g], [h], [i]]` corresponds to the payoff matrix:

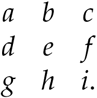

For this example, we set *a* = 0, *b* = – 1, *c* = 3, and so forth.

~~~
In [2]: payoff_entries = [[0], [-1], [3], [-1], [0], [1], [3], [1], [0]]
        simplex = plot_static(payoff_entries, background=False, edge_eq=True)
~~~

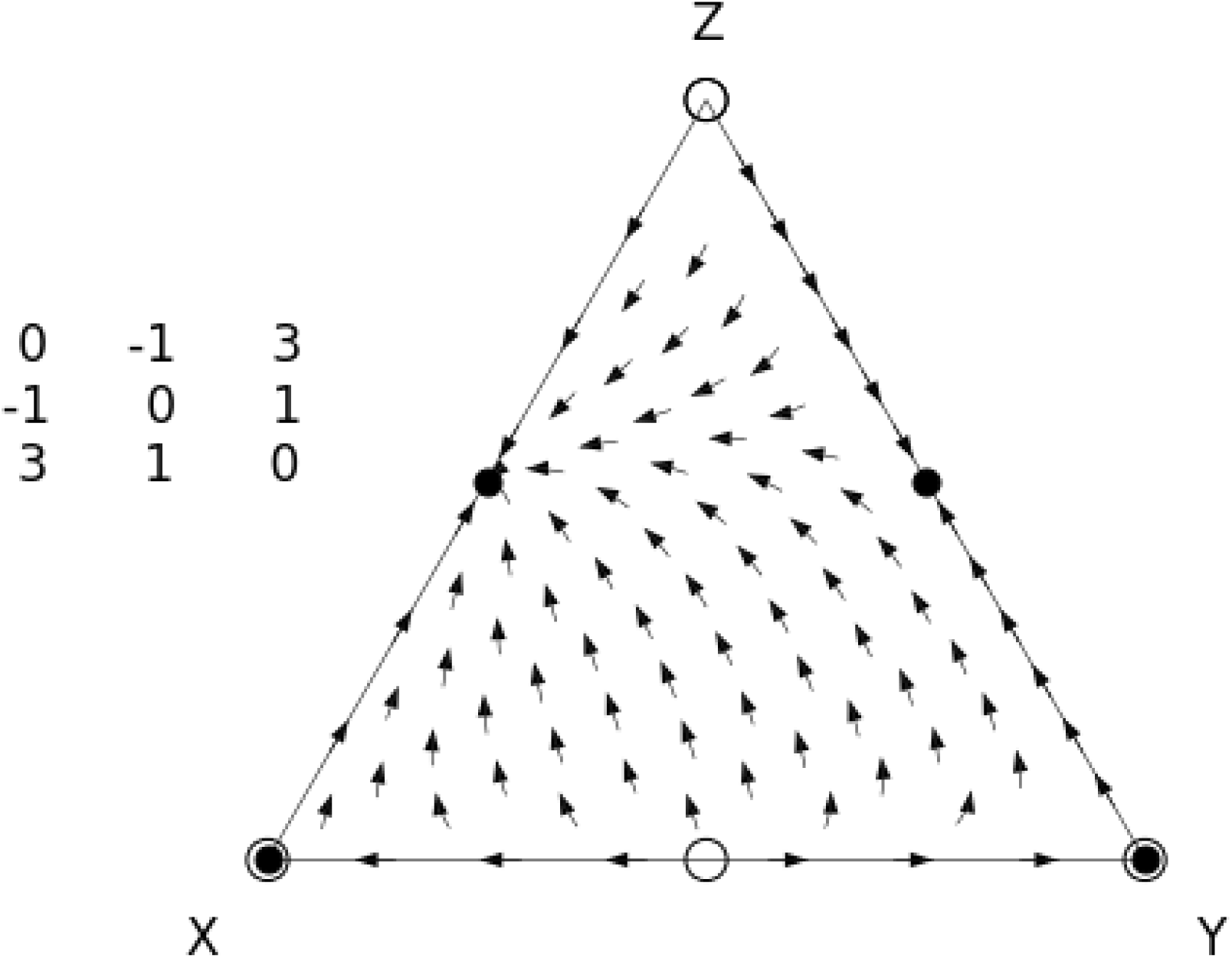

The plot that is produced very much resembles that of Bomze. We use the same notation of stable and unstable equilibria as filled circles and unfilled circles, respectively. However, at this time, the package is only capable of determining whether equilibria are stable for initial conditions exactly on that edge. A point that begins precisely on the right edge will travel to the stable equilibrium approximately midway between the vertices labeled “Y” and “Z” and will then stay there. If that point is off that edge even very slightly, that stable equilibrium will instead act as a saddle point, first attracting the point, and then repelling it towards the globally-stable equilibrium on the left edge.

### 1.3 A Further Example

While there are many use cases for this software, our group’s particular focus is on the mathematical modeling of cancer and cancer therapies through evolutionary game theory [7, 8, 9]. For details on analytical treatments of evolutionary games, please see Artem Kaznatcheev’s blog Theory, Evolution, and Games Group.

We demonstrate the features of `egtplot` via an example drawn from work modelling the interactions between cancer and healthy cells. Consider the following game from [10] which describes the evolutionary game between three strategies of cells, labeled “S” (Stroma), “D” (micro-nenvironmentally dependent), and “I” (microenvironmentally independent):

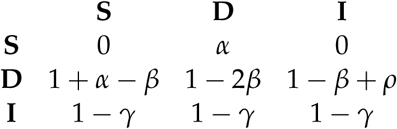

where *α* is the benefit derived from the cooperation between a S cell and a D cell, *γ* is the cost of being microenvironmentally independent, *β* is the cost of extracting resources from the microenvironment, and *ρ* is the benefit derived by a D cell from paracrine growth factors produced by I cells. This paper studies how the healthy cells that make up the majority of the prostate can cooperate and compete with mutant protate cells to produce a clinically-detectable prostate cancer.

Using our package, we can quickly and easily analyze this game numerically and visually. To start, let us choose some simple values for each parameter: *α* = 1, *β* = 1, *γ* = 1, and *ρ* = 1.

We first create a helper function to create our list of entries to the payoff matrix for our chosen values of *α*, *β*, *γ*, and *ρ*.

~~~
In [3]: def get_payoff(alpha, beta, gamma, rho):
            return [[0, alpha, 0],
                    [1 + alpha - beta, 1 - 2 ∗ beta, 1 - beta + rho],
                    [1 - gamma, 1 - gamma, 1 - gamma]]
~~~

Now that we have the entries of our payoff matrix, we can pass them to `plot_static` to view the resulting simplex. Here, we call `plot_static` with the default plotting arguments, but use our helper function and label the vertices appropriately.

~~~
In [4]: parameter_values = [[1], [1], [1], [1]]
        labels = [‘S’, ‘D’;, ‘I’]
        simplex = plot_static(parameter_values, custom_func=get_payoff, vert_labels=labels)
~~~

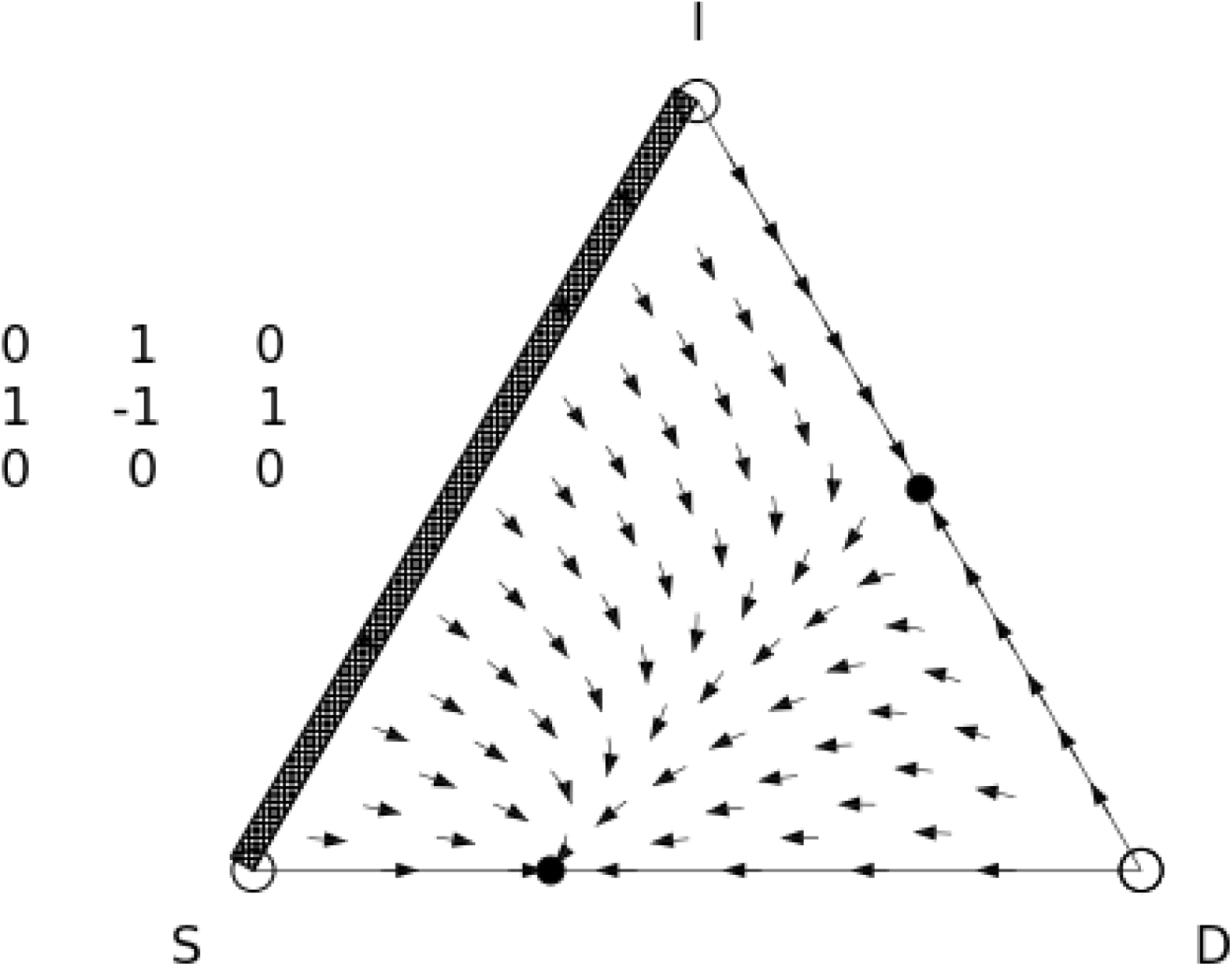

We see that there is a saddle point on the D-I edge and a globally-stable equilibrium on the S-D edge. Additionally, the hacthed rectangle on the S-I edge indicates that every point on that edge is an equilibrium‐‐i.e. no matter where it is on the edge, a point will not move. Finally, we see unstable equilirbia at all three vertices.

## 2 Function Arguments

Though the default arguments create a visually appealing simplex plot, they may not be appropriate for every situation. Our function `plot_static` takes a number of arguments that allow for extensive customization.

The arguments that `plot_static` takes are:

- `payoff_entries` This is the list of lists of entries of the payoff matrix that we described above.
- `custom_func` This is the helper function that can be designated to turn a functional form of the payoff matrix (as in our example above) into the list of lists of payoffs. Defaults to None.
- `generations` This is the number of epochs to simulate forward the ODEs. More generations means more total time that is simulated. Defaults to 6.
- `steps` This is the number of steps simulated per generation. More steps means a denser simulation. Defaults to 200.
- `background` Set this argument to True if the background of the simplex should be colored according to the speed at which the strategy mix is changing at each point. Set this to False for a blank background. Defaults to `False`.
- `ic_type` This argument controls the placement of initial conditions to be simulated within the simplex. It has three possible values: ‘`grid`’, which places initial conditions on a grid ‘`edge`’, which places initial conditions evenly along each edge, and ‘`random`’, which distributes initial conditions randomly throughout the simplex. In the case of ‘`grid`’ and ‘`random`’, 11 initial conditions are placed on each edge. This is necessary so that the background speed coloring, if present, covers the entirety of the simplex. Defaults to `grid`.
- `ic_num` For `ic_type = ‘edge’` or `ic_type = ‘random’`, this sets the number of initial conditions on each edge or in the interior, respectively. Defaults to 100.
- `ic_dist` When `ic_type = ‘random’`, this sets the minimum distance initial conditions can be from each other, to keep them from being too dense. Defaults to 0.05.
- `ic_color` This sets the color of the markers for the initial conditions. It is passed directly to the matplotlib plotting functions, so can accept any color that matplotlib itself does. Defaults to black.
- `paths` This argument controls whether the paths of each initial condition are plotted. When this is `True`, the arrows seen above are replaced with dots. Defaults to `False`.
- `path_color` This sets the colormap used in the plotting of the paths. It is passed directly to the matplotlib plotting functions, so can accept any colormap that matplotlib itself does. Defaults to inferno.
- `edge_eq` The function has the ability to compute the equilibria of the three subgames that form the edges of the simplex. If this argument is set to `True`, these equilibria are displayed. The types of displayed equilibria are: - cross-hatched rectangles which indicate that every point on that edge is an equilibrium, - open circles which indicate that point is an unstable equilirbium, - closed black circles which indicate that point is a stable equilibrium, and - closed grey circles which indicate that point is semistable (has a second-derivative which is zero). Defaults to `True`.
- `display_parameters` This argument controls whether the payoff matrix used to construct the simplex is displayed above the simplex. Defaults to True.
- `vert_labels` This argument accepts a list to label the vertices of the simplex, beginning with the bottom left vertex and proceeding anti-clockwise. Defaults to [`‘X’, ‘Y’, ‘Z’`].

## 3 Illustrative Examples

Here are some examples of our same payoff matrix passed to `plot_static`, but with changes to the default values. First, we change to displaying the paths taken by each initial condition, which may provide a more granular view of the dynamics within the simplex or if there is a particular initial condition or set of initial conditions that are particularly interesting.

~~~
In [5]: simplex = plot_static(parameter_values, custom_func=get_payoff, vert_labels=labels, paths=True)
~~~

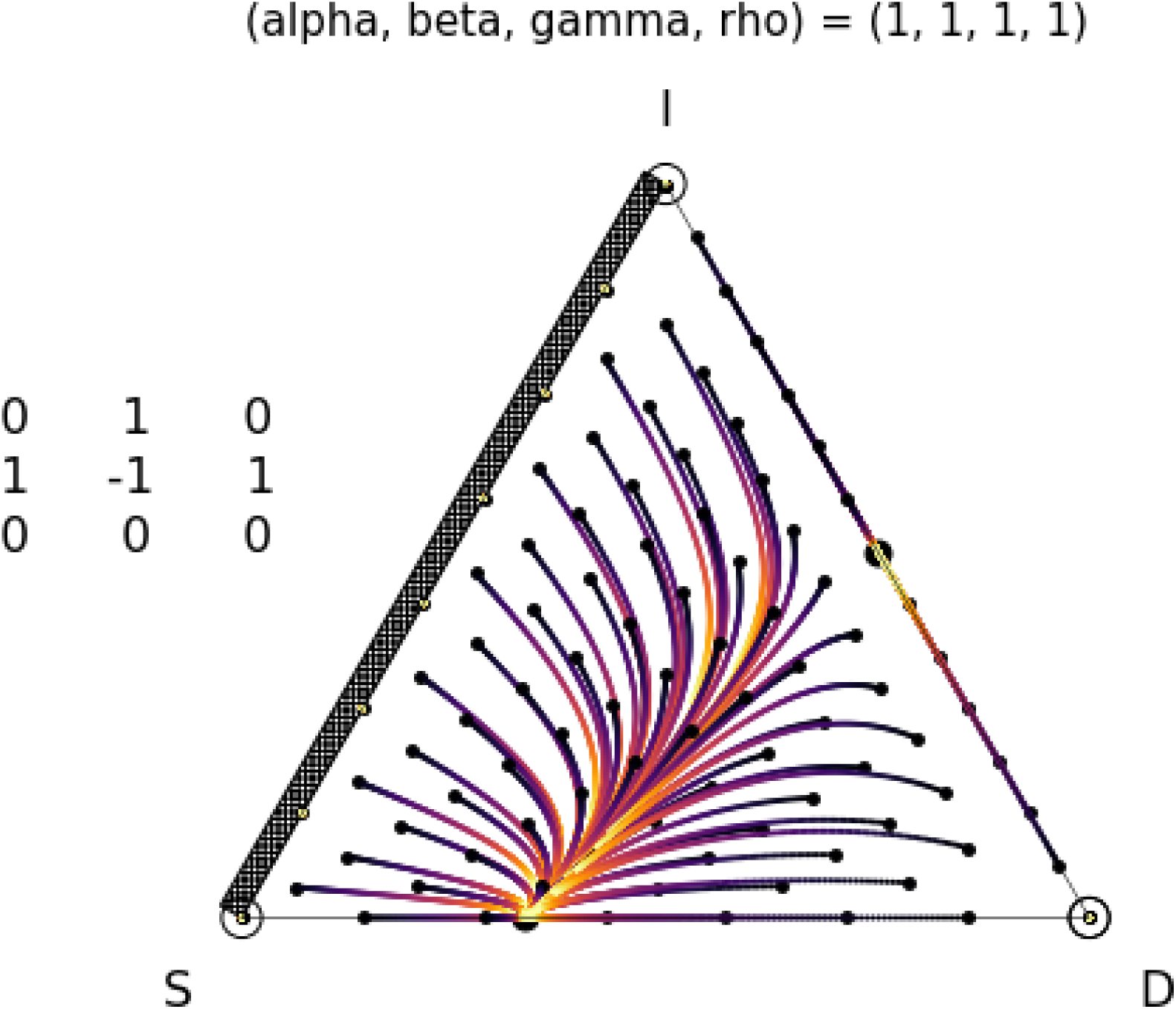

Next, we color the background of the simplex by the speed at which the points would travel along their trajectories. This also displays a colorbar to the left of the plot with arbitrary units. Note that the coloring does not influence the direction of travel in any way, but that stable equilibria are always located in an area of locally low speed.

~~~
In [6]: simplex = plot_static(parameter_values, custom_func=get_payoff, vert_labels=labels, background=True)
~~~

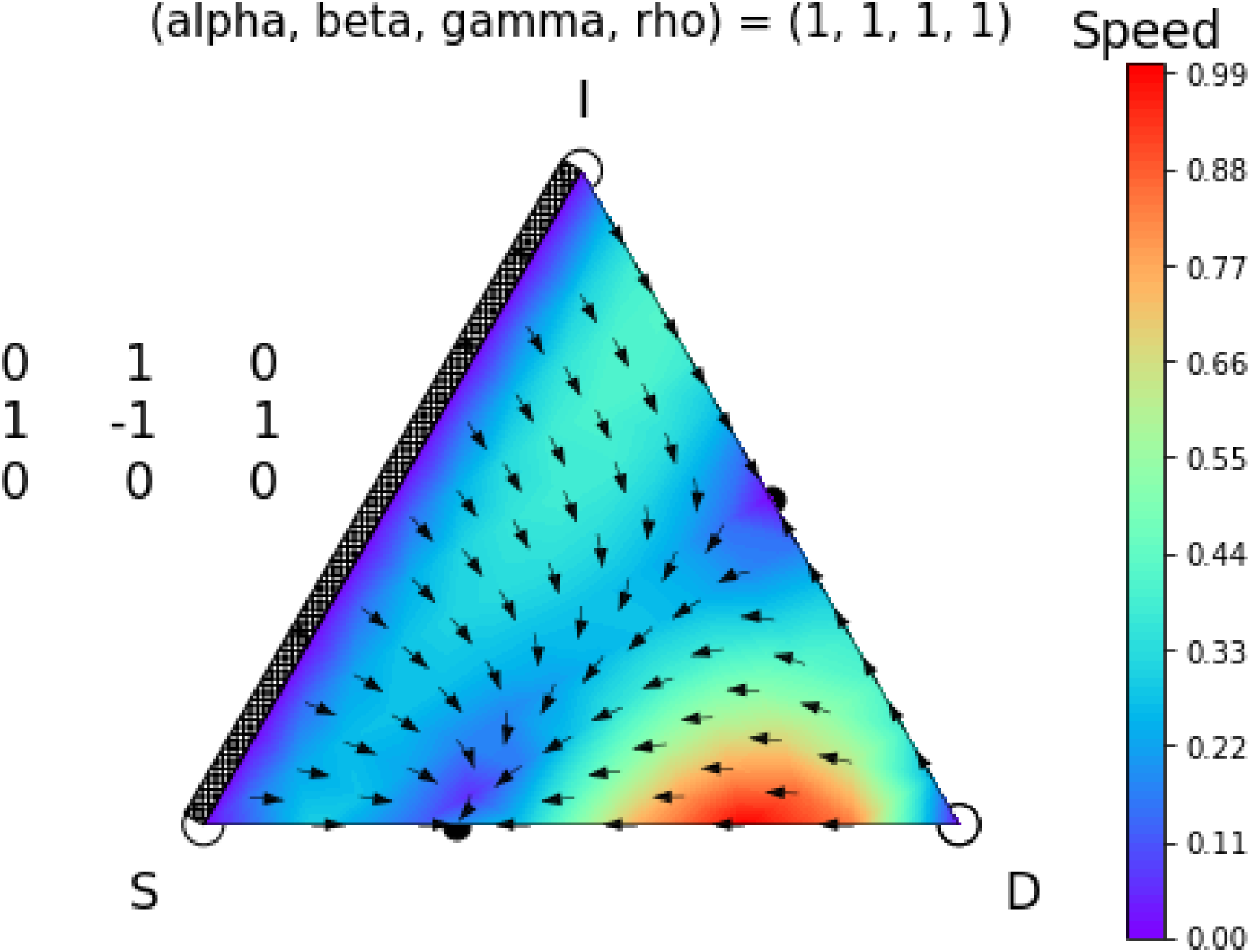

Finally, we again display paths, but change the colormap, increase the number of steps and generations to make the paths longer and smoother, and use a random distribution of initial conditions within the simplex. We also do not display the equilibria.

~~~
In [7]: simplex = plot_static(parameter_values, custom_func=get_payoff, vert_labels=labels,
                                paths=True,
                                generations=10,
                                steps=2000,
                                ic_type=‘random’,
                                path_color=‘viridis’,
                                edge_eq=False)
~~~

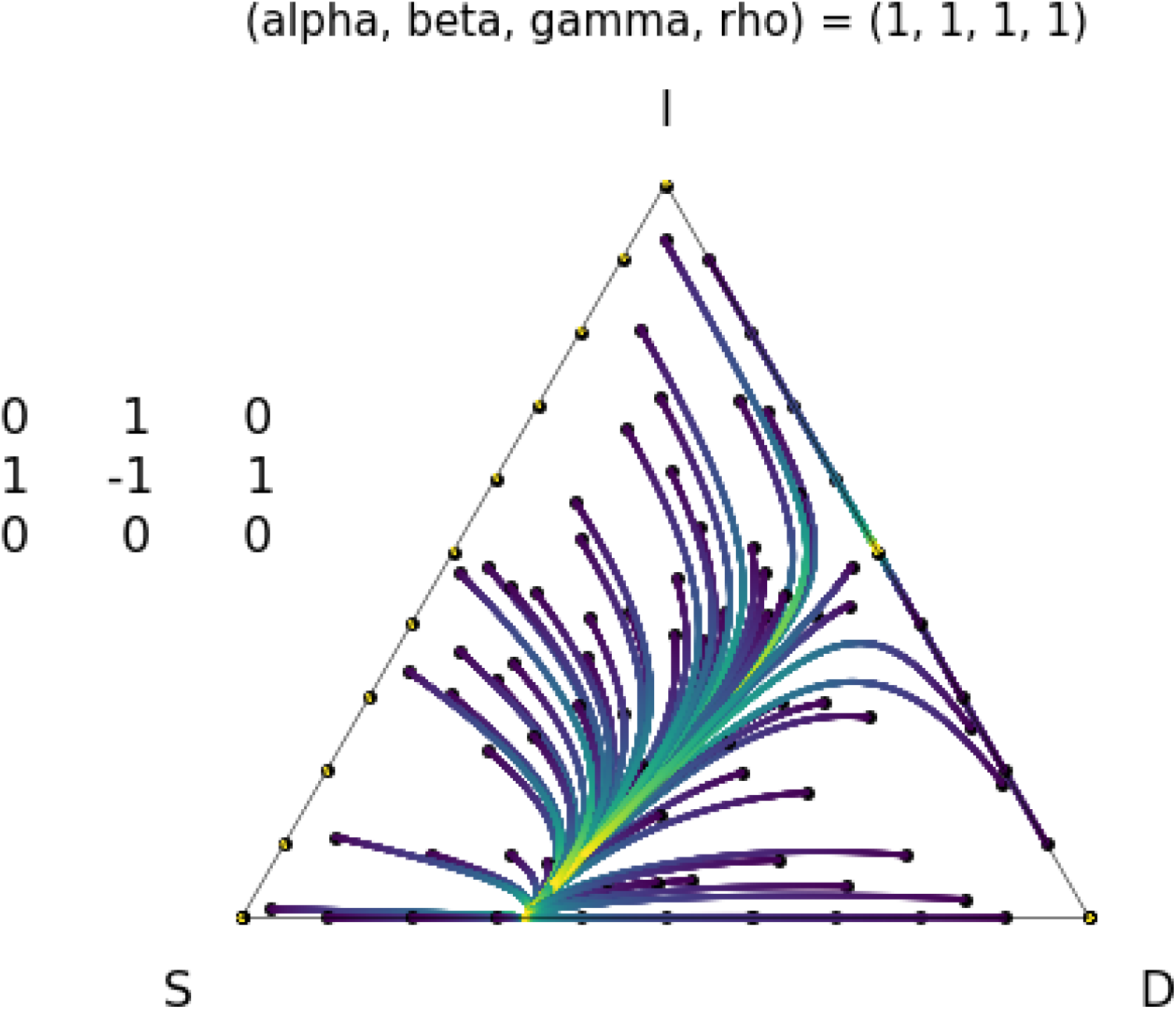

## 4 Parameter Sweeps

The function `plot_static` also has the ability to handle mulitple values for entries in the payoff matrix. In our example above, we may want to test two different values for *α*, 1 and 2 to see how they would independently affect the dynamics of the game.

To do this, we simply add an entry to the corresponding element of our list of parameter values. The lists within our parameter value list can be any length, as long as there is one list for each parameter. We continue to use our helper function, `get_payoff` defined above, to parse these parameter values into a list of nine lists.

~~~
In [8]: parameter_values = [[1, 2], [1], [1], [1]]
        simplex = plot_static(parameter_values, custom_func=get_payoff, vert_labels=labels)
~~~

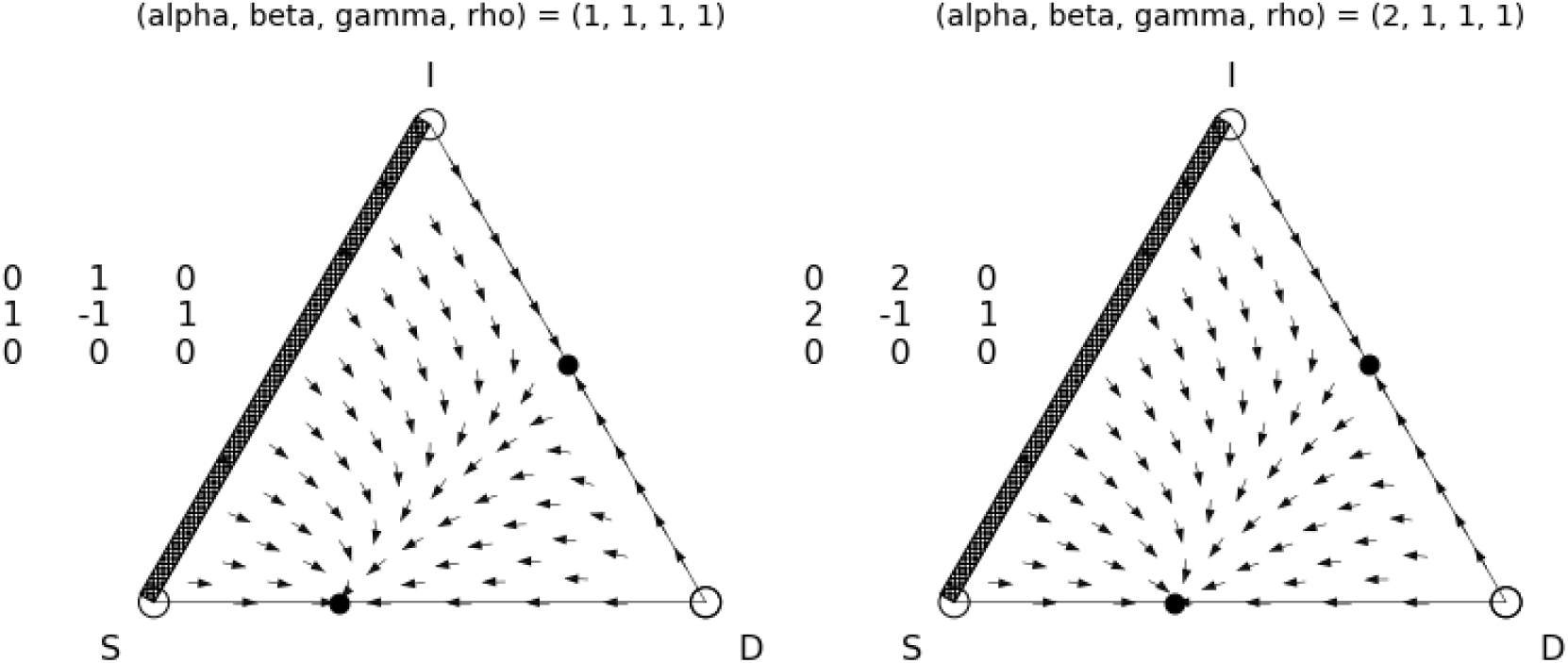

We can extend our parameter sweep to the other parameters, as well. In this example, in addition to testing *α* equal to 1 and 2, we also test*β* equal to 1 and 3. We also display the speeds within the simplex.

~~~
In [9]: parameter_values = [[1, 2], [1, 3], [1], [1]]
        simplex = plot_static(parameter_values, custom_func=get_payoff, vert_labels=labels, background=True)
~~~

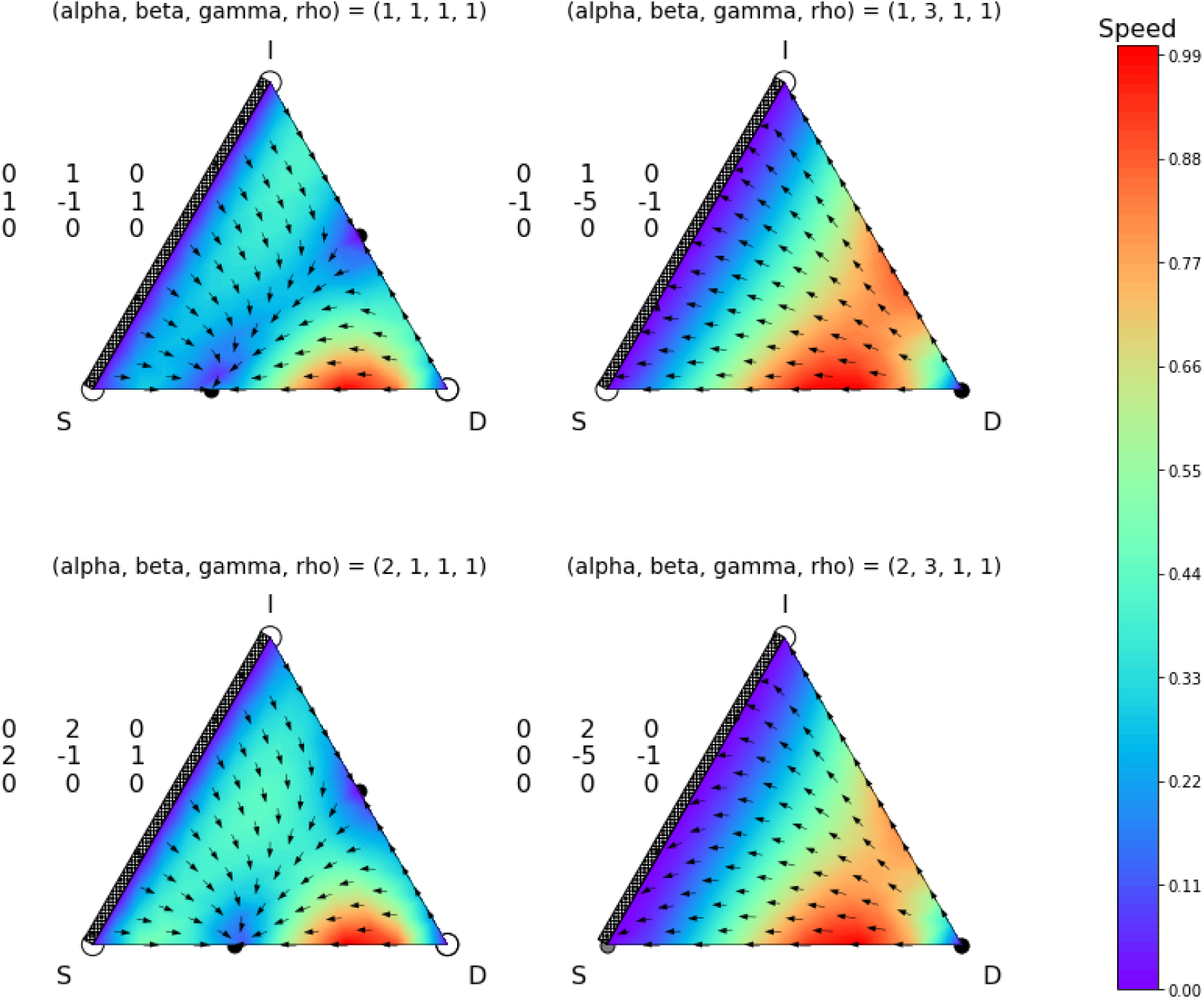

When only two parameters are varied at a time, `plot_static` increases the first varying parameter along the y-axis of the subplots and the second along the x-axis, as seen above. When more than two parameters are varied, the subplots are fit into the smallest square that will fit all of the subplots, but the y-axis and x-axis structure when varying only two parameters is lost.

Below, we vary *α*, *β*, and γ, display the paths taken by each initial condition, and choose 20 initial conditions that are equally distributed along each edge.

~~~
In [10]: parameter_values = [[1, 2], [1, 3], [1, 4], [1]]
         simplex = plot_static(parameter_values, custom_func=get_payoff, vert_labels=labels,
                                 steps=600,
                                 paths=True,
                                 ic_type=‘edge’,
                                 ic_num=20)
~~~

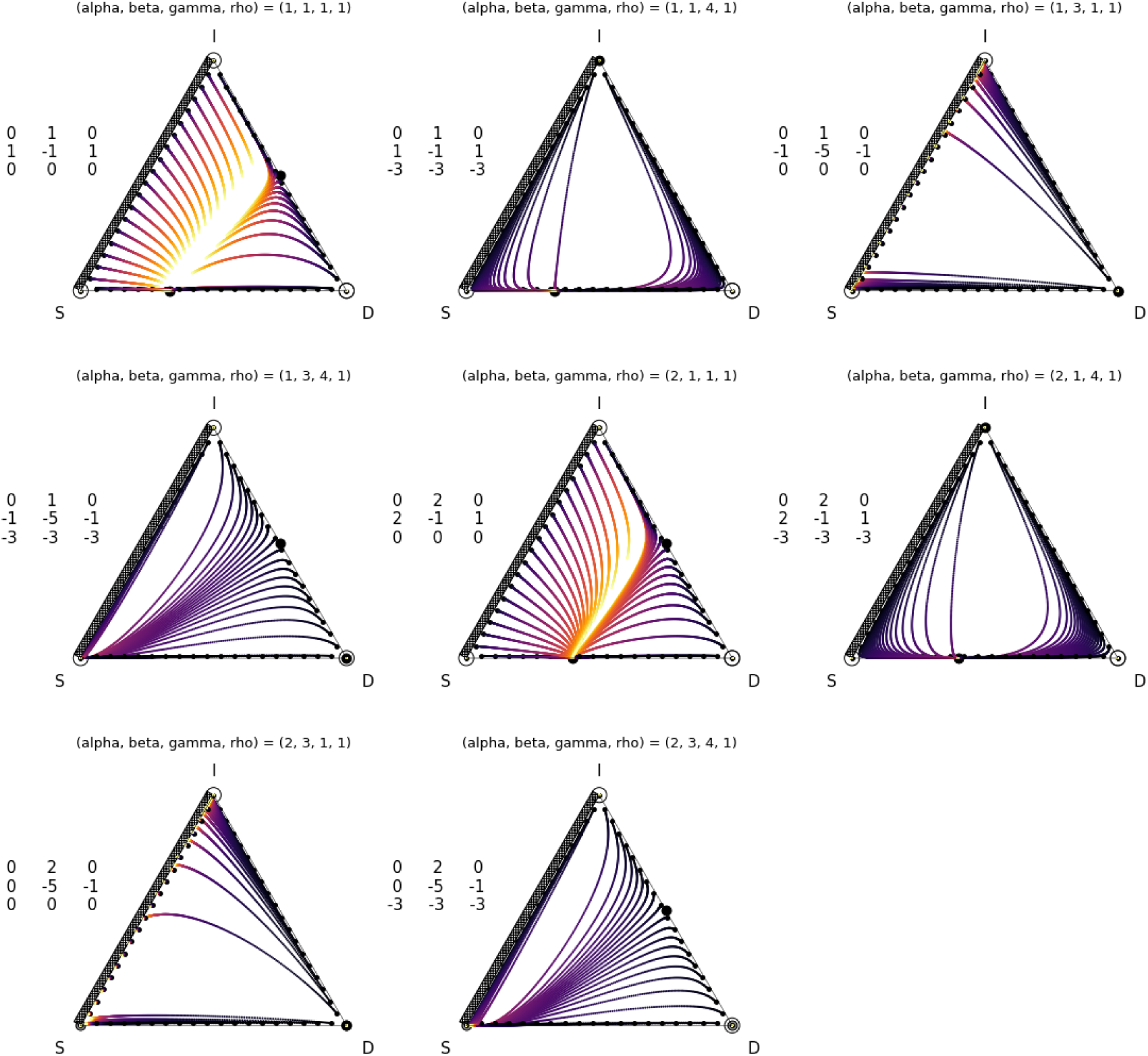

All arguments that work on individual plots work for subplots of parameter sweeps as well. The package does not currently have the functionality to assign different arguments such as `ic_type` or `path_color` to individual subplots.

~~~
In [11]: parameter_values = [[1, 2], [1, 3], [1, 4], [1]]
         simplex = plot_static(parameter_values, custom_func=get_payoff, vert_labels=labels,
                                 steps=2000,
                                 background=False,
                                 ic_type=‘random’,
                                 paths=True,
                                 path_color=‘hot’,
                                 edge_eq=False)
~~~

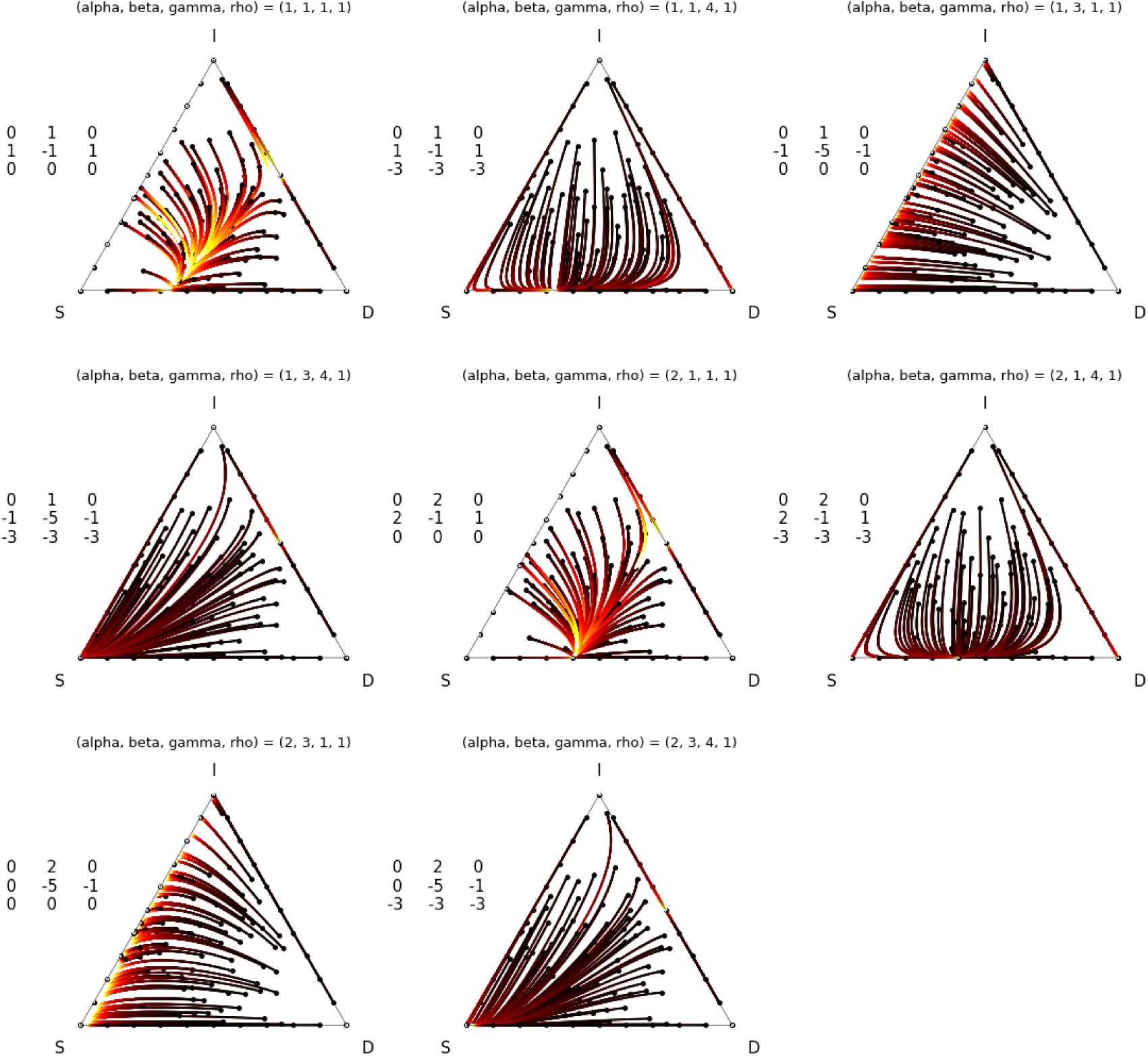

## 5 Animations

Our software package also allows for the creation of animated simplices through the use of the `plot_animated` function. Because we cannot embed the animations in this document, we have omitted much of the description of this functionality here. Please see our github repository for complete description and demonstration jupyter notebook.

Many arguments are shared between plot_static and `plot_animated`. These are `payoff_entries`, `custom_func`, `vert_labels`, `generations`, `ic_type`, `ic_num`, `ic_dist`, `edge_eq`, and `display_paramters`. The defaults for `generations` and `steps` have been changed to 30 and 120, respectively. There are, however, also arguments that `plot_animated` does not accept. These are `background` and `paths`.

Arguments specific to `plot_animated` are:

- `num_fps` This argument specifies an integer number of frames per second for the animation. The duration of the animation is equal to `steps` divided by `num_fps` in seconds. Defaults to `30`.
- `dot_color` This argument takes a string which sets the color of the moving dots. If ‘`rgb`’ is specified, the dots will be colored with red, green, and blue values corresponding to the population’s initial proportion of each species (i.e., a population that is monotypic X will be depicted as a dot that is entirely blue while a population that is an equal mix of Y and Z will be 50% green and 50% red). Defaults to ‘`rgb`’.
- `dot_size` This sets the size of the moving dots. Defaults to `20`.

As with its static counterpart, `plot_animated` has the ability to plot parameter sweeps. The sweeps are passed and plotted in an identical manner to the static examples above.

## 6 Acknowledgements

- This material is based upon work supported by the National Science Foundation under Agreement No. 0931642 (Mathematical Biosciences Institute at Ohio State University).
- We gratefully acknowledge the work of Hanna Schenk whose code on her [github](https://github.com/HannaSchenk/RQchaos) inspired this project.

